# Adaptive genomic variation associated with environmental gradients along a latitudinal cline in *Rana temporaria*

**DOI:** 10.1101/427872

**Authors:** Alexandra Jansen van Rensburg, Maria Cortazar-Chinarro, Annsi Laurila, Josh Van Buskirk

**Affiliations:** Department of Evolutionary Biology and Environmental Studies, University of Zurich, Switzerland; School of Biological Sciences, University of Bristol, UK; Department of Ecology and Genetics, Uppsala University, Sweden

## Abstract

*Rana temporaria* occur across a large geographic and environmental gradient in Scandinavia. Several studies involving common garden experiments have established adaptive divergence across the gradient. The main objective of this study was to determine the extent of neutral and adaptive genetic divergence across the latitudinal gradient. Here we sequence genome-wide markers for 15 populations from six regions sampled from southern Sweden to Finland. Using a multivariate approach we find that 68% of the genomic variation is associated with climate or geographically structured climate. Using outlier scans and environmental association analyses we identify a set of potentially adaptive loci and examine their change in allele frequency associated with different climatic variables. Using a gradient forest analysis we identify points along three of the climate variables where allele frequencies change more rapidly than expected if it were a linear association. We identify a large threshold effect associated with BIO5 (mean temperature during the warmest month) which is seen as a rapid change in southern Sweden. By comparing the change in neutral and adaptive allele frequencies across the whole gradient, we identify southern Sweden as a region with the largest divergence between the datasets. This suggests small changes in the climate may result in a mismatch between the adaptive genotypes and the environment in these populations. Overall this study shows that genomic analyses can provide a powerful complement to common garden experiments to improve our understanding of adaptive divergence across heterogeneous landscapes.

## Introduction

The geographic distribution of genetic variation across a species range is an important determinant of population persistence under changing environmental conditions (Hoffmann & Sgrò 2011). To mitigate future biodiversity loss due to climate change it is important that we identify the most important environmental drivers of adaptive divergence and determine how genetic variation contributes to adaptation. Landscape genomics methods have been employed to identify potentially adaptive loci across many study systems (reviewed in Rellstab *et al*. 2015). However, adaptation across environmental gradients are often characterised by divergence in multiple phenotypic traits with polygenic genetic underpinnings (Pritchard & Di Rienzo 2010; Yeaman 2015). This presents a challenge when examining non-model organisms, where the lack of genomic resources is often prohibitive in determining the genomic architecture and genetic basis of adaptation (Manel *et al*. 2016). However, by developing our understanding of the distribution of genetic variation (adaptive and neutral) in geographic and climate space, we can draw meaningful conclusions about the ecological determinants of species distributions and adaptive divergence without pinpointing the underlying causal variants (Jones *et al*. 2013; Fitzpatrick & Keller 2015; Forester *et al*. 2016).

Two main statistical models have been developed to identify potentially adaptive loci while accounting for population structure (reviewed in Hoban *et al*. 2016). Population genetic methods identify loci with higher than expected differentiation (usually measured by F_ST_) compared to neutral expectations based on population structure (Luikart *et al*. 2003). These methods are effective for detecting selective sweeps associated with strong selection acting on a few beneficial loci. However, the signal of divergence is difficult to detect if there is high gene flow between populations adapted to different conditions, or when the adaptive traits are polygenic with many small changes in allele frequency additively contributing to adaptation (Kawecki & Ebert 2004; Pritchard & Di Rienzo 2010). The second kind of model, termed Environmental Association Analysis (EAA; reviewed in Rellstab *et al*. 2015), is aimed at finding associations between allele frequencies and environmental variables, thus does not rely on strong sweeps underlying adaptation. This approach is particularly useful for comparing the importance of different environmental variables and for mapping spatial changes in allele frequencies of adaptive loci. However, EAA assumes a linear relationship between allele frequencies and environmental gradients (Thomassen *et al*. 2010; Fitzpatrick *et al*. 2015), thus confining inferences to linear responses. But non-linear patterns of genetic variation along environmental gradients are probably common, judging from laboratory studies of organismal physiology (Angilletta’s 2009 thermal adaptation book, for example), and could be important for identifying populations or geographic regions that are especially vulnerable to climate change. Thus, it is important to modify EAA such that non-linear relationships between allele frequencies and environmental gradients can be detected in nature. Here we combine a modelling approach to loci identified using F_ST_ outlier and EAA methods to identify such non-linear associations with environment in the European common frog, *Rana temporaria* across a latitudinal gradient.

*Rana temporaria* is widespread across Europe and occur throughout Scandinavia (Sillero *et al*. 2014). Populations occur across a wide range of habitats, which suggests adaptive divergence across the species range. Common garden experiments across 1600-km of the Scandinavian latitudinal gradient have confirmed this by establishing extensive latitudinal variation in larval and adult life history traits (Merila *et al*. 2000; Laugen *et al*. 2003b, 2005a; Palo *et al*. 2003a; Lindgren & Laurila 2005). The geographic scale of the gradient provides an interesting system to investigate the genomics of adaptation to environment, because there is little gene flow between populations adapted to different environments.

The main objective of this study is to determine the extent of neutral and adaptive genetic divergence across the latitudinal gradient. Specifically, we aim to 1) characterise the neutral genetic structure, 2) determine the proportion of the genome associated with environmentally driven adaptive divergence, and 3) identify environmental thresholds to adaptation by examining the non-linear response of adaptive loci to climate variables. We were particularly interested in determining whether there are non-linear relationships between adaptive allele frequencies and environmental variables. Such a response would suggest that there is a threshold along that particular environmental variable that requires a larger change in allele frequency than would be expected based on the gradual change in that environmental variable. The results have important implications for identifying populations and geographic regions that would be particularly vulnerable to changing environments.

## Methods

### Sampling & DNA extraction

To determine intra-specific population structure and adaptive variation, 163 individuals were collected from six geographic regions across ~1500 km of the Scandinavian latitudinal gradient (Fig. 1; Table 1). Each region was represented by one to three populations (where a population is a pond), for a total of 15 populations in the final dataset. At each sampling site, we sampled approximately 10 eggs from 20-30 freshly -laid clutches (less than two days old). The eggs were transported to the laboratory at Uppsala University where they were raised in separate containers kept in climate room at 16 °C. Tadpoles were raised to Gosner stage 25 (Gosner 1960), whereafter they were euthanized with an overdose of MS222, preserved in 96% ethanol and stored at 4 °C until DNA extraction. Total DNA was extracted from one individual per clutch (henceforth family) using the Qiagen DNeay blood and tissue kit (Qiagen, CA, USA).

**Figure 1.**
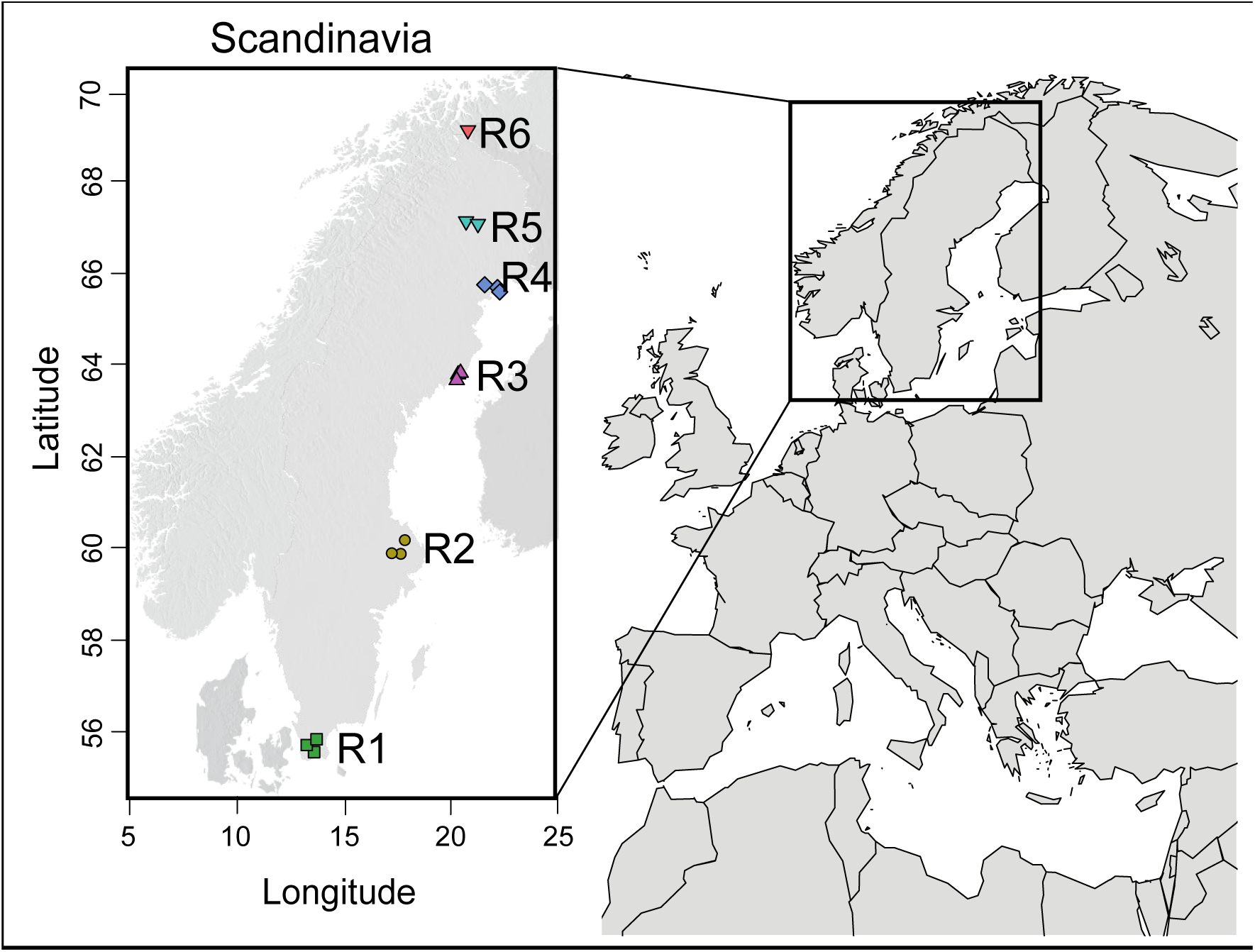
Sampled populations across the Fennoscandian latitudinal gradient. One to three populations were sampled within each of six regions. Symbols represent the five genetic clusters found at K=5, the most likely number of divisions found by DAPC, PCAdapt, and SNMF analyses. Geographic regions are named R1 - R6 from south to north along the latitudinal gradient.

**Table 1.**
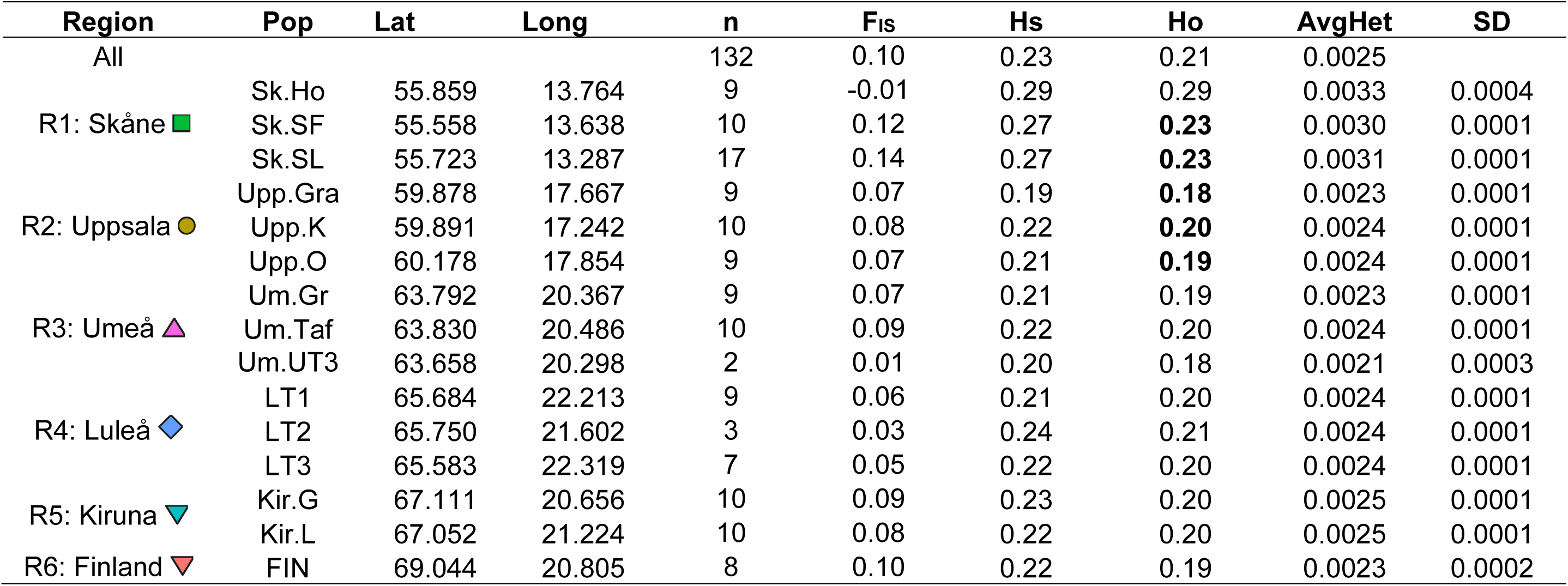
Summary of the diversity statistics calculated per population. The geographic coordinates (Lat, Long) are shown for each population (Pop). The geographic region (Region) to which the populations belong is shown along with the abbreviation (R1-R6) used throughout the manuscript. The number of individuals included in the analyses (n) are shown. Gene diversities (Hs), deviation from random random mating (F_IS_), observed heterozygosity (Ho), and the average number of heterozygous sites across all sequenced sites (AvgHet) with standard deviation (SD) are reported per population.

| Region | Pop | Lat | Long | n | F<sub>IS</sub> | Hs | H<sub>O</sub> | AvgHet | SD |
| --- | --- | --- | --- | --- | --- | --- | --- | --- | --- |
| All |  |  |  | 132 | 0.10 | 0.23 | 0.21 | 0.0025 |  |
| R1: Skåne | Sk.Ho | 55.859 | 13.764 | 9 | -0.01 | 0.29 | 0.29 | 0.0033 | 0.0004 |
|  | Sk.SF | 55.558 | 13.638 | 10 | 0.12 | 0.27 | **0.23** | 0.0030 | 0.0001 |
|  | Sk.SL | 55.723 | 13.287 | 17 | 0.14 | 0.27 | **0.23** | 0.0031 | 0.0001 |
| R2: Uppsala | Upp.Gra | 59.878 | 17.667 | 9 | 0.07 | 0.19 | **0.18** | 0.0023 | 0.0001 |
|  | Upp.K | 59.891 | 17.242 | 10 | 0.08 | 0.22 | **0.20** | 0.0024 | 0.0001 |
|  | Upp.O | 60.178 | 17.854 | 9 | 0.07 | 0.21 | **0.19** | 0.0024 | 0.0001 |
| R3: Umeå | Um.Gr | 63.792 | 20.367 | 9 | 0.07 | 0.21 | 0.19 | 0.0023 | 0.0001 |
|  | Um.Taf | 63.830 | 20.486 | 10 | 0.09 | 0.22 | 0.20 | 0.0024 | 0.0001 |
|  | Um.UT3 | 63.658 | 20.298 | 2 | 0.01 | 0.20 | 0.18 | 0.0021 | 0.0003 |
| R4: Luleå | LT1 | 65.684 | 22.213 | 9 | 0.06 | 0.21 | 0.20 | 0.0024 | 0.0001 |
|  | LT2 | 65.750 | 21.602 | 3 | 0.03 | 0.24 | 0.21 | 0.0024 | 0.0001 |
|  | LT3 | 65.583 | 22.319 | 7 | 0.05 | 0.22 | 0.20 | 0.0024 | 0.0001 |
| R5: Kiruna | Kir.G | 67.111 | 20.656 | 10 | 0.09 | 0.23 | 0.20 | 0.0025 | 0.0001 |
|  | Kir.L | 67.052 | 21.224 | 10 | 0.08 | 0.22 | 0.20 | 0.0025 | 0.0001 |
| R6: Finland | FIN | 69.044 | 20.805 | 8 | 0.10 | 0.22 | 0.19 | 0.0023 | 0.0002 |

### ddRAD sequencing, de novo assembly, and variant calling

To establish a genome-wide marker set, double digest restriction-site-associated DNA sequencing libraries were prepared with the restriction enzymes *EcoRI* and *MseI*, using a modification of the protocol by Peterson *et al*. (2012). We constructed 6 libraries comprising 48 samples each for single-end sequencing (125bp) on an Illumina HiSeq 2500 v4 at the Functional Genomics Center, University of Zurich. Individual samples were identified by unique 5-bp barcodes. Raw sequence reads were demultiplexed using the process_radtags package from Stacks (Catchen *et al*. 2011). Demultiplexing was based on a unique 9-bp sequence for each individual (5-bp unique barcode + 4-bp restriction enzyme recognition site), with 1 mismatch allowed. Adapter and other Illumina-specific sequences were removed using Trimmomatic v0.33 (Bolger *et al*. 2014).

*De novo* assembly and variant calling was implemented using pyRAD (Eaton 2014) which first finds clusters within individuals based on a clustering threshold and minimum depth, and then clusters these loci between individuals (using the same clustering threshold), and identifies loci found in a user specified number of individuals. Clustering thresholds of 90-99% sequence similarity were tested; an optimum of 94% was chosen because it maximised nucleotide diversity and minimised the estimated number of paralogs in the dataset. A maximum of 4 sites with a Phred quality score <20 were allowed per sequence. Clusters were kept if they had >5x coverage per individual and were found in at least 4 individuals.

### SNP validation

The putative variants identified though the pyRAD pipeline were filtered for possible sequencing errors, paralogs, and uninformative SNPs. The following filters were applied: 1) SNPs that were genotyped in less than 50% of individuals were removed using the *--max-missing* function in VCFtools v0.1.12b (Danecek *et al*. 2011). 2) Loci with a minor allele frequency less than 0.05 in the full dataset were removed as they are more likely to be sequencing error, and if they are true variants they are uninformative and likely to bias tests for selection (Roesti *et al*. 2012). 3) Using a sliding-window of 10-bp we tested whether the number of variants increased towards the end of the sequence. No significant difference was found between bins, thus the sequences were not trimmed further. 4) We reduced linkage in the dataset we included only one variant per locus using the *--thin* function in VCFtools. 5) We assessed whether loci were in Hardy-Weinberg equilibrium (HWE) within each population using the *--hardy* function in *PLINK* (Purcell *et al*. 2007). Loci with an observed heterozygosity more than 0.5, and loci that deviated significantly from HWE based on the exact test (p<0.05; (Wigginton *et al*. 2005) were removed from the dataset if they occurred in more than 5 populations. 6) We then calculated linkage disequilibrium for each locus pair per population in PLINK v 1.07 (Purcell *et al*. 2007). Loci with *r^2^* > 0.8 never occurred in more than 5 populations, so no loci were excluded at this step. 7) Finally we excluded individuals with more than 55% missingness.

### Summary statistics

Nucleotide diversity was calculated for each sample as the frequency of heterozygous bases (IUPAC codes) from the *pyRAD* output, and means were calculated per population. Although these calculations are based on sequences before final filters are implemented, two reasons convince us that these results are robust: 1) *pyRAD* calculates a binomial probability that a base is homozygous or heterozygous based on a maximum likelihood approach of jointly estimating the heterozygosity and sequencing error rate from the base frequencies within each individual’s stacks. If the read depth of the stack falls below a user-set threshold, or is too low to make a statistical base call, the base remains undetermined. Thus the final base calls per individual should be fairly robust (Eaton 2014). 2) The post-pyRAD filters outlined in the previous paragraph do not address sequencing or SNP-calling errors, but rather minimises missingness, systematic biases, and linkage between loci.

Further summary statistics were calculated in R v3.3.1 (R core team 2016) with the *hierfstat v0.04-22* and *adegenet v2.0.1* packages (Goudet 2005; Jombart 2008; Jombart & Ahmed 2011). Gene diversities (Hs), deviation from random mating (F_IS_), and observed and expected heterozygosity are reported per population (Table 1).

### Population structure

Pairwise population differentiation was estimated using Weir and Cockerham’s F_ST_ (Weir & Cockerham 1984) as implemented in *hierfstat* (Goudet 2005). We visualised the genetic distance between populations with a principal component analysis (PCA) implemented in *PCAdapt v3.0.3* (Luu *et al*. 2016). The following analyses were conducted in *adegenet*. To test for isolation by distance, we calculated the correlation between log transformed pairwise geographic distances and scaled pairwise genetic distances (F_ST_/1-F_ST_) (Rousset 1997). We tested for significance using a Mantel test. We quantified the proportion of genetic variation that explained differentiation within and between populations with an analysis of molecular variance (AMOVA). Finally, we performed a discriminant analysis of principal components (DAPC) to determine the most likely number of clusters in the dataset and visualise broad-scale population structure.

### Climate Data

We obtained climate data from WorldClim v1.4 (Hijmans *et al*. 2005) at a resolution of 2.5 minutes of degrees using the R package *raster v.2.5-8* (Hijmans *et al*. 2015). Many of these variables are derived from the same data and are highly correlated. To reduce the redundancy in the climate variables retained for analyses, we first calculated correlation between all variables, and then removed variables if they exceeded a correlation threshold. We calculated the absolute values of pairwise ranked correlation (Spearman’s rho) between all 19 BioClim variables from WorldClim, longitude, latitude, and season length. Season length was calculated as the number of days above 6 and 8 °C at each sampling site, since 6 °C is approximately the development threshold of *R. temporaria* tadpoles (Laugen *et al*. 2003a). We reduced redundancy in the environmental dataset by detecting pairs of variables that had an absolute correlation >0.8, and then eliminating the one that had the highest mean correlation with all other variables (Kuhn et al. 2016). This procedure retained five BioClim variables, and these were used as the environmental variables in all remaining analyses (Fig. 2).

**Figure 2.**
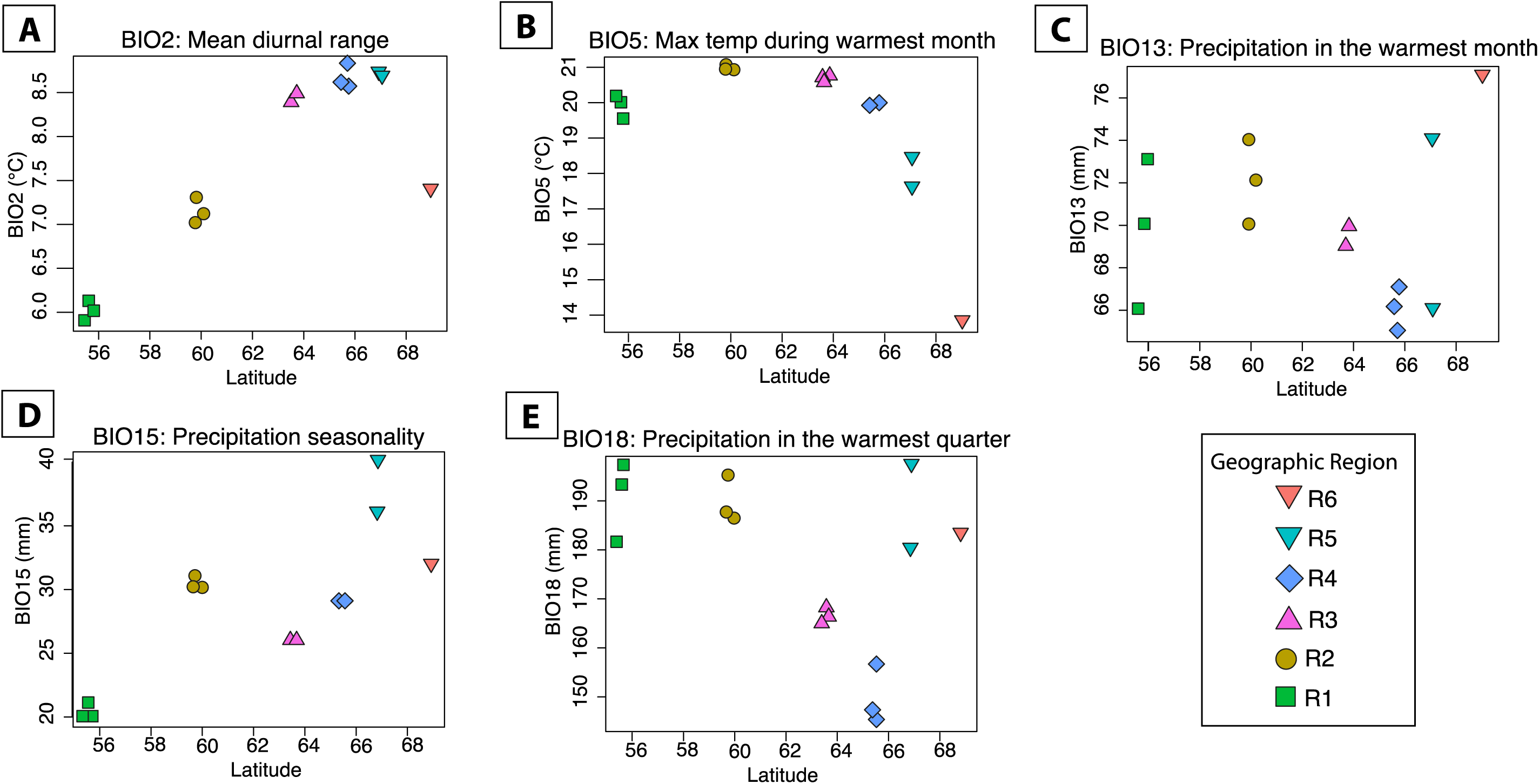
Latitudinal distribution of the five BioClim variables used in this study. Each point represents a sampled population.

### Relative contribution of environment and IBD to genomic differentiation

We investigated the effects of climate and geography on neutral genetic structure using full and partial redundancy analyses (RDA) with variance partitioning. Redundancy analysis is a form of multivariate regression, which can be used when both the predictor and response variables are multivariate (Legendre & Legendre 2012). As a canonical extension of multiple linear regression, RDA identifies a set of orthogonal linear predictor variables that explains the most variation in a set of linear response variables. In this case each RDA axis represents a set of co-varying loci (response variables), which are correlated with co-varying environmental variables (predictor variables). RDA has greater power to detect multivariate genotype-environment relationships than methods based on distance matrices or Mantel tests (Legendre & Fortin 2010).

We created two matrices as response variables: 1) the 5 climate variables identified above, centered and standardised, and 2) geographic coordinates of each sampling site. The response matrix was the minor allele frequencies of 2081 loci for each individual. We ran a sequence of nested models to partition variation in climate and geography as explanatory variables of allele frequencies: (1) the full model including both climate and geography; (2) climate only, with the influence of geography partialled out (climate | geography); and (3) geography only, with climate partialled out (geography | climate). The difference in the variance explained by model (1) minus the sum of models (2) and (3) was interpreted as the contribution of climate and geography acting together. Overall and residual variance was calculated for each model, and the model significance was tested with 999 permutations. The RDA was conducted using the R package *vegan v2.4-1* (Oksanen *et al*. 2015).

### Signatures of adaptation: Outlier detection and genotype-environment associations

We created two datasets containing loci potentially under selection using two common approaches: 1) Outlier analyses to identify loci that are more differentiated between populations than expected under a neutral model, and 2) Environmental Association Analyses (EAA) to identify loci strongly associated with an environmental variable (reviewed in Rellstab *et al*. 2015). These approaches are likely to identify different loci, since their underlying assumptions are different. The Outlier dataset comprised loci identified with *PCAdapt* (Luu *et al*. 2016), and from the X_T_X statistic calculated in *bayenv2* (Günther & Coop 2013). The EAA dataset comprised loci identified using *bayenv2* and LFMM (Frichot *et al*. 2013).

*PCAdapt* identifies outlier loci as those that are more associated with population structure than expected. We used the R package *pcadapt 3.0.4* (Luu *et al*. 2016), which calculates a vector of z-scores of the how related each SNP is to the first K principal components, where K is the user-specified number of population clusters. A Mahalanobis distance is then calculated for each SNP to determine whether it deviates from the main distribution of z-scores. These scores are scaled by a constant, the genomic inflation factor, which produces a chi-squared distribution of values with K degrees of freedom. K was calculated as the most likely number of genetic clusters after testing K 1-20 and inspecting the scree plot of the proportion of explained variance for each K (Fig. S1). Based on these results, we chose K=5 for further analyses. We used a false discovery rate of 10% to identify outlier loci.

*Bayenv2* estimates genotype-environment associations and an F_ST_-like statistic (X_T_X) while correcting for covariance of allele frequency between populations due to neutral processes. We used *bayenv2* to identify loci for both datasets. First we estimated the neutral covariance matrix based on 500 randomly selected loci. Two independent runs with 100000 MCMC iterations were run. We tested for convergence within each run by calculating Pearson’s product-moment correlation (*cor.test* in R) between the final matrix and nine matrices printed out at 10000 step intervals (9 correlations). We constructed distance-based trees to determine whether relationships among populations remained constant within and between runs. Convergence between runs was calculated as the correlation between the final matrix in each run. Since these were highly correlated, we arbitrarily chose the final matrix from the first run as our final covariance matrix. The full model was run using this covariance matrix, a file containing standardised measures of each environmental variable, and a genotype file containing SNP counts across all populations. We ran a non-parametric test that calculates the Bayes factor, Spearman’s *p*, and Pearson’s correlation coefficient for each genotype-environment association (-t -c -r). In addition we calculated the X_T_X population differentiation statistic (-X). For this test, Gunther *et al*. (2013) suggest ranking loci by their X_T_X statistic rather than selecting those above a specific threshold.

We conducted three independent runs with *bayenv2* of 100,000 MCMC iterations each for each of the five genotype-environment associations, and tested convergence by calculating the correlation between runs for each statistic (BF, *p*, X_T_X). We also compared the overlap in loci identified in the top 5%, 6-10%, and 11-15% ranked loci based on the X_T_X statistic for each environmental variable to ensure the repeatability of the results. We then ran an additional 7 independent *bayenv2* runs, and calculated the median result across all 10 runs as our final output. We selected the top 100 ranked loci based on the X_T_X statistic for the outlier dataset. For the EAA dataset we selected loci with a log10 Bayes Factor (BF) >0.5 (Kass & Raftery 1995), and absolute Spearman’s rho (p) >0.3.

Finally, we screened the dataset for additional EAA loci using the Latent Factor Mixed-effect Model (LFMM; Frichot *et al*. 2013; Frichot & François 2015). LFMM calculates the correlation between genotype and environment while simultaneously accounting for population structure with latent factors incorporated in the model. The number of latent factors is user specified, and should represent the number of genetic clusters (K) that best describes the population structure in the dataset. As suggested by Frichot & François (2015), we estimated the most likely K by evaluating the cross-entropy criterion for K1-10 using the function *snmf* in the R package *LEA*. The most likely K was 5, which is consistent with the population structure analyses described above, and therefore LFMM was run with K=5 for each of the five environmental variables. The Gibbs sampler algorithm was run five times for each environmental variable, with 10000 cycles and a burn-in of 5000 cycles. The median of the resulting correlation scores (z-scores) was calculated across all five runs. The authors suggest a recalibration of the mean z-scores by lambda; that is the square of the mean z-scores divided by the median of a chi-squared distribution with one degree of freedom (λ; ~0.455). Lambda should be close to one - but more importantly, this should produce the correct adjusted p-value frequency distribution. Adjustment of λ can correct for liberal or conservative p-value distributions. We evaluated the effect of λ = 0.45-1.00 on the p-value distribution for each of the 5 environment-genotype associations. The shape of the distribution did not change much, but the frequency of p-values >0.1 increased as λ increased (i.e. the correction was more conservative). Thus, for a lambda close to one and the correct adjusted p-value distribution, we chose λ = 0.85 for BIO2, BIO5, BIO15, and BIO18, and λ = 0.45 for BIO13. To control for false discoveries, we applied a Benjamin-Hochberg adjustment with a false discovery rate of 5%.

### Genomic Turnover across ecological gradients

We assessed how genomic variation changes across Scandinavia and whether important climatic thresholds occur by fitting a Gradient Forest model (Ellis *et al*. 2012) to each of the SNP datasets. We compare the change in allele frequency between the adaptive loci (Outlier and EAA dataset) and a Reference dataset composed of all the remaining loci. The Gradient forest model was developed as an extension of the random forest model to assess community level responses to ecological gradients, and has recently been applied to genomic data to detect non-linear change in allele frequencies along ecological gradients (Fitzpatrick & Keller 2015). It is a machine-learning ensemble approach that fits multiple regression trees between allele frequency and environmental variables. A set of decision trees is built to describe change in allele frequency across the predictor variable range. Each split is determined by minimising the "impurity" in the data, i.e. minimising the sums of squares of the allele frequency and thus maximising the tree fit. This means the split will always describe the biggest change in allele frequency at the current point in the tree, and a relative split importance can be calculated.

The response variable was the minor allele frequencies (MAF) for each SNP dataset. We included only loci variable in more than 4 populations to ensure robust regressions. The predictor variables were the 5 BioClim variables described above. To account for unsampled geographic structure in the dataset, we also included Moran’s Eigenvector Map variables (MEM; Dray et al. 2006), which are orthogonal vectors maximising spatial autocorrelation between sampled locations. Broad-scale spatial structure is most likely explained by the most positive eigenvectors (Manel et al. 2010b, 2012; Sork et al. 2013), so we included the first half of the positive MEMs here; in total three MEM vectors.

The gradient forest model was fit to each SNP dataset using the R package *gradientForest* (Ellis *et al*. 2012). We constructed 2000 regression trees per SNP, with default values for the variable correlation threshold (0.5), the number of candidate predictor variables sampled at each split (2), and the proportion of samples used for training (~66%) and testing (~33%) each tree. The relative importance (R^2^) of each predictor variable was calculated as the weighted mean of the proportion of variance explained by the validation data. The cumulative importance for the change in allele frequency for each locus was calculated as the sum of the split importance across climatic variables, and the mean allelic turnover per climatic variable was calculated for each of the three SNP datasets.

Changes in allele frequency across the landscape were visualised by transforming each climatic variable by the genomic importance calculated for each SNP dataset; i.e. we produced a transformed dataset for the Reference, EAA, and Outlier datasets. The three transformed datasets were produced for climate data extracted from a raster stack covering eastern Sweden and northernmost Finland. For each dataset, the transformed variables were reduced by PCA, and a colour from the RGB colour palette in R was assigned to each of the first three principal components. Thus, for each geographic point along the latitudinal gradient, a single colour was used to represent the genomic composition in three-dimensional principal component space. To compare the genomic turnover between the neutral and two candidate SNP datasets, we calculated the distance in genomic space at each geographic point as the Procrustes residuals between the pairs of transformed matrices calculated above. The genomic difference between datasets was normalised and mapped in geographic space as before.

## Results

### ddRAD data generation and variant filtering

The final dataset consisted of 132 individuals from 15 populations (2-17 individuals per population; Table 1). There were 2081 SNP loci, with a mean depth of 17.9 - 25.5x and genotyping rate of 72-97% per population. A summary of the number of raw reads, the output from pyRAD, and the final dataset can be found in Table S1.

### Genetic variation and Population structure

The highest genetic diversity was found in the southernmost region, R1, while the rest of the gradient was characterised by lower genetic diversity that was similar across all regions (Table 1). Specifically, the mean gene diversity (H_S_) and heterozygosity (H_E_) in Skåne were 0.28 and 0.25, respectively, while they were only 0.22 and 0.19 across the rest of the gradient. Similarly, the frequency of heterozygous sites across all sequenced sites averaged ~1/300 (0.0031) in R1, and ~1/400 (0.0024) across the rest of the gradient.

Measures of genetic differentiation showed evidence of population structure within and between regions. AMOVA indicated that most genetic variation (68.72%) was found within populations, while significant variation was found among populations within regions (6.05%) and among (25.23%) regions (Table S2). The mean global F_ST_ was 0.21 (Fig. 3, Table S3), suggesting strong population structure on average between populations. However, pairwise genetic distance was much lower within (F_ST_ = 0.06) than between (F_ST_ = 0.16) regions, and there was significant isolation by distance (R=0.434, p=0.001) across the sampled area.

**Figure 3.**
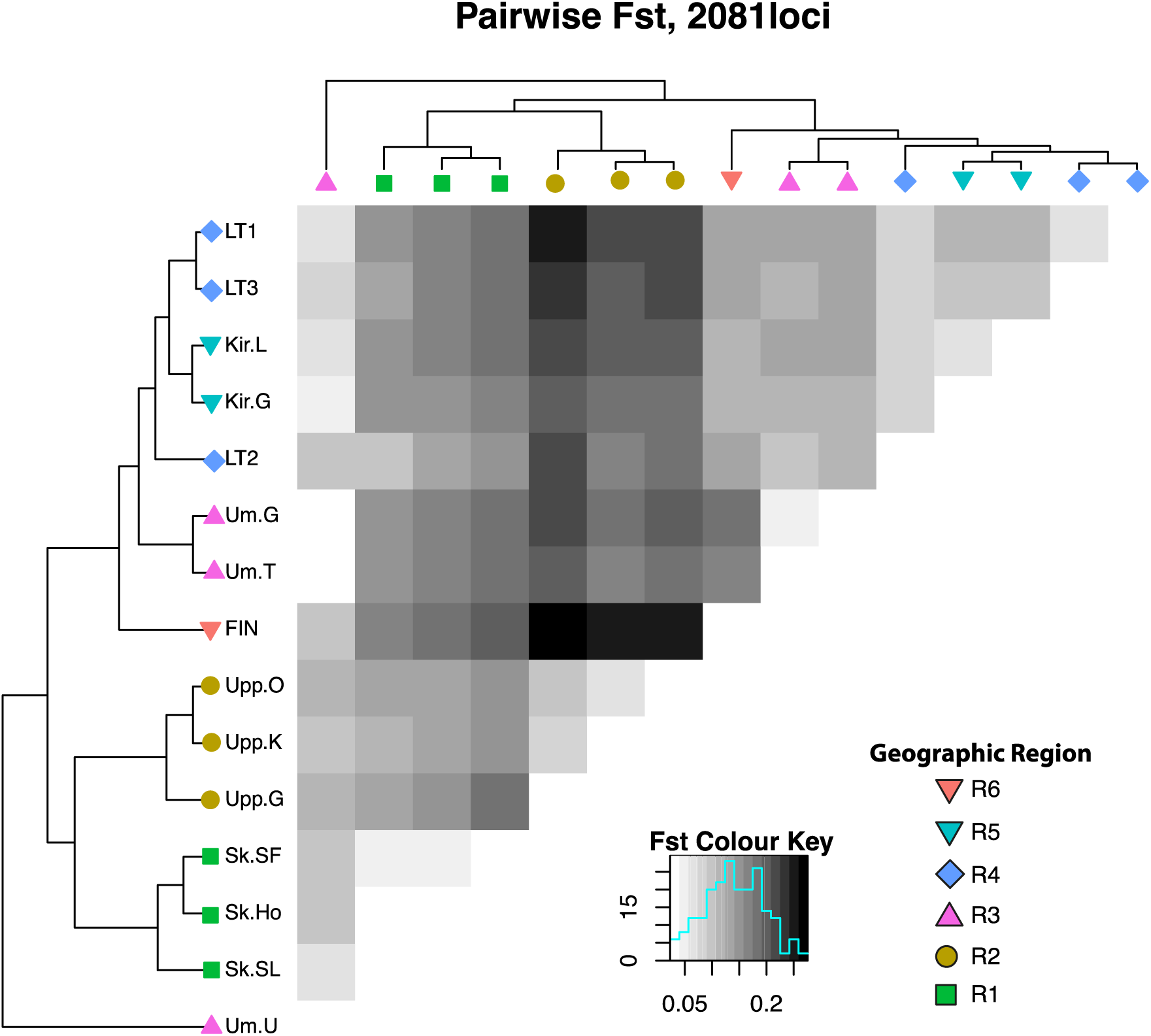
Heatmap of the pairwise genetic divergence (F_ST_) between all sampled populations. Darker squares indicates higher F_ST_, with colour scaled as shown in the key. Geographic regions are differentiated with shapes corresponding to populations in Fig. 1. Colours correspond to the PCA shown in Fig. 4. The dendrogram shows the population structure between southern (R1-R2) and northern (R3-R6) Scandinavian populations.

To determine the broad-scale population structure, we first visualised the genetic distance between populations using PCA, and then estimated the most likely number of genetic clusters using DAPC. The first two axes of the PCA explained approximately 24% of the variance. PC1 (~15%) partitioned the two southern regions (R1 and R2) from the rest, and PC2 (~9%) partitioned populations latitudinally, with a graded differentiation from R2 to R6 (Fig. 4).

**Figure 4.**
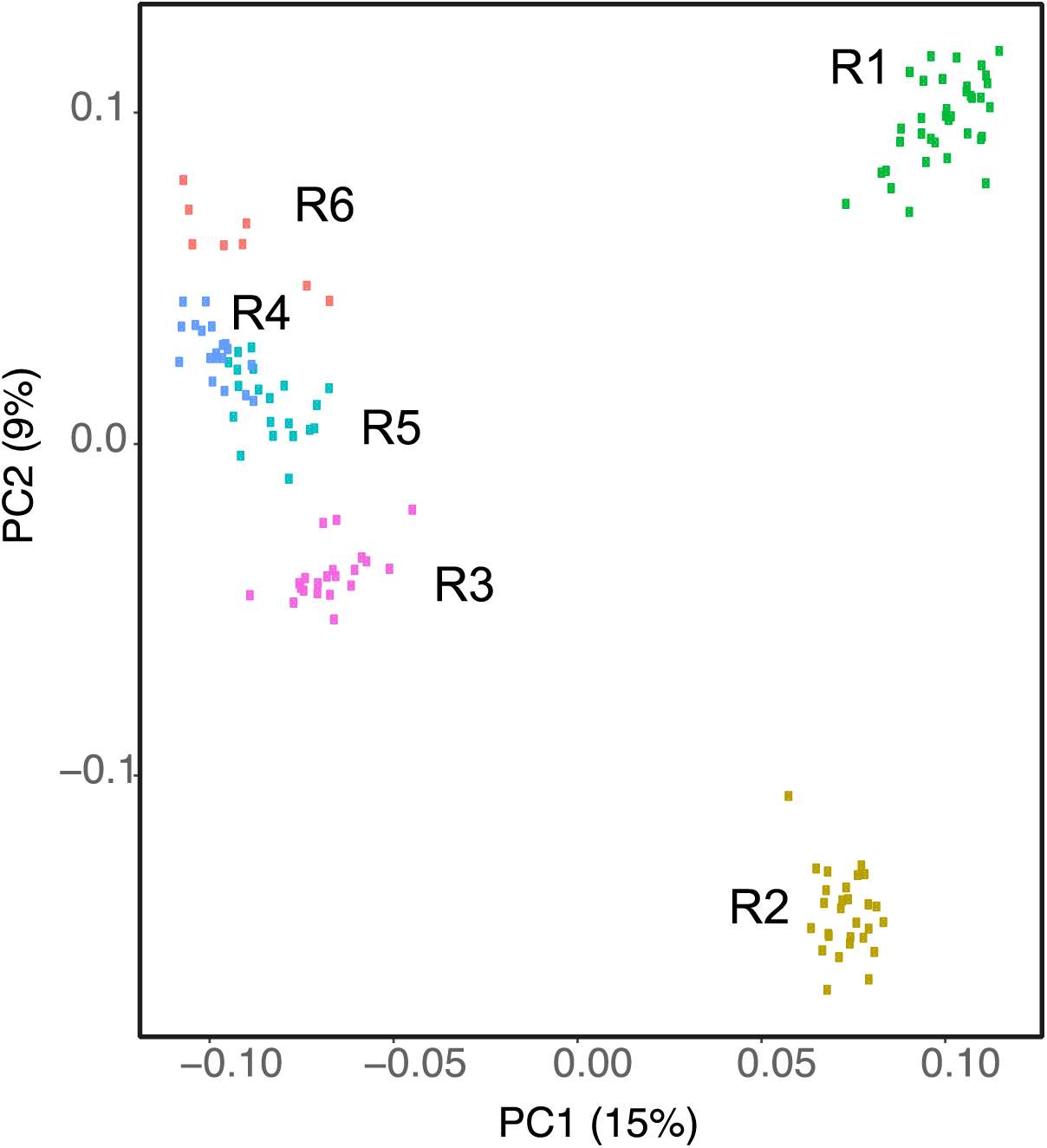
Graph of the first two axes of a Principal Component Analysis of all populations. The proportion variation explained by each axis is shown in brackets. Regions are shown in different colours.

Discriminant analysis of principal components (DAPC) predicted five genetic clusters, corresponding to R1, R2, R3, R4, and the three northern populations (Fig. 1). When separating the dataset into increasing numbers of clusters, populations grouped out sequentially from south to north, except that R5 and R6 always grouped together.

### Redundancy Analysis

The full RDA model included climate and geographic coordinates and explained 76.6% of the total genetic variation (*p=0.001*). Based on the partial RDA, climate was significantly associated with genetic variation (climate | geography; *p=0.002*), and explained 49.5% of the total variation. The variation explained by geography alone (geography | climate) was much less (11.4%) and was non-significant. The proportion of genetic variation explained by spatially structured climate (climate n geography) was 39.1%.

### Signature of adaptation: Outlier detection and genotype-environment associations

*PCAdapt* identified 50 outlier SNP loci and the X_T_X statistic from *bayenv2* returned the top 100 outlier loci. A total of 28 loci were identified by both methods, so that the Outlier dataset comprised 122 unique loci (Fig. S2).

The EAA dataset comprised loci identified using *Bayenv2* and LFMM as associated with the five chosen BioClim variables. *Bayenv2* identified 123 unique loci (6% of the total loci tested), with 13% of these loci associated with multiple environmental variables (Fig. S3). LFMM identified 398 unique loci (~19% of the total loci), with ~30% associated with multiple environmental variables (Fig. S4). Only 22 loci were identified by both LFMM and *Bayenv2;* thus, the final EAA dataset comprised 499 unique loci (Fig. S5).

There were 56 loci present in both the Outlier and EAA datasets (Fig. S6). This represents 45.9% of the Outlier dataset, and 11.2% of the EAA dataset.

### Genomic Turnover

We used the mean R^2^ averaged over loci in each dataset as a measure of the fit importance and thus how informative each dataset was (Table 2). The Reference and EAA datasets performed similarly, with mean R^2^ values of 37.6% and 38.9%, respectively. The Outlier dataset showed a better fit on average, with a mean R^2^ of 56.5%. The frequency distributions of R^2^ values differed among datasets. The Reference and EAA datasets both had fairly flat distributions, with 30.5% and 34% of loci with R^2^>0.5. The R^2^ of the Outlier dataset was right skewed, with 67% of loci identified with R^2^>0.5.

**Table 2.**
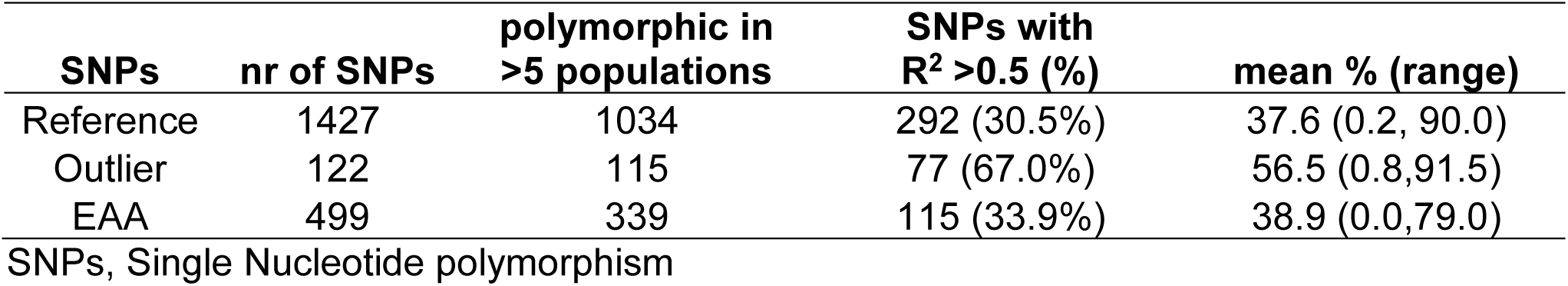
Summary of the three SNP datasets and their results from the Gradient Forest analysis. The model was fit only to SNPs that were variable in more than five populations.

The most important variables for all three datasets were distance and either MEM1 or MEM2 (Fig. 5). This suggests that geographic distance is an important determinant of the genomic differences between populations, and that the MEM variables captured important environmental variation that had not been included in the model. When considering only the BioClim variables, BIO5 and BIO2 were the most important variables for the Outlier dataset, and BIO5 for the EAA dataset. Notably, the Outlier dataset had a higher fit importance (R^2^) to the BIO2 and BIO5 variables than the EAA or Reference dataset. This suggests that these variables explain the change in allele frequency in the Outlier dataset better than the change in allele frequency in the Reference our EAA datasets.

**Figure 5.**
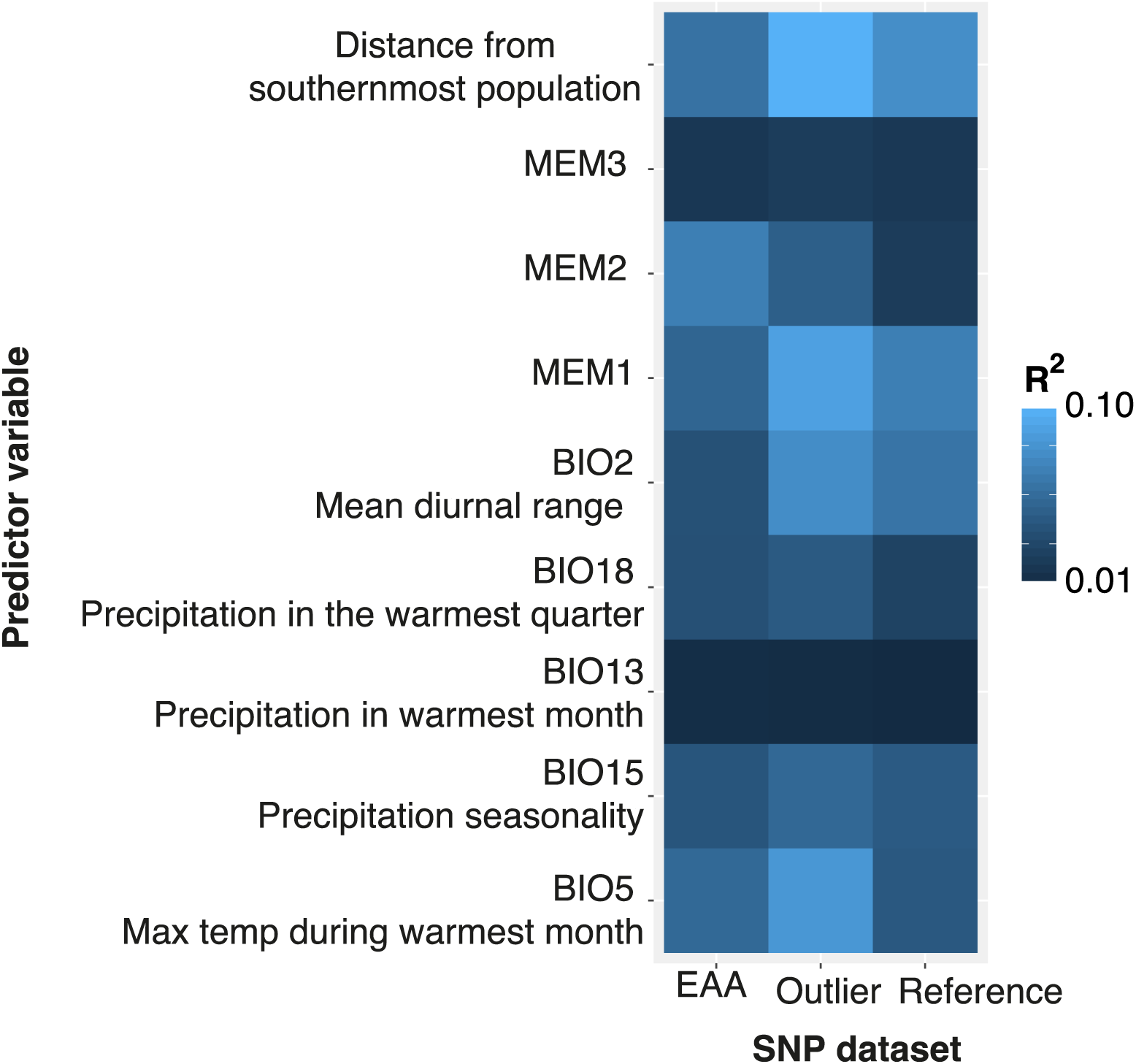
The relative importance (R2) of the predictor variables used in the Gradient Forest analysis, calculated as the weighted mean proportion of variance in allele frequency explained by a given environmental variable. Results are shown for the five BioClim variables, geographic distance, and three Moran’s Eigenvectors (MEM1-3) explaining geographic structure in the dataset. The columns show results for the SNP datasets. EAA: Adaptive loci from environmental association analyses; Outlier: Adaptive loci identified using PCAdapt and XTX statistic in BayEnv2; Reference: the remaining loci. Lighter colours indicating higher importance; e.g. BIO2 and BIO5 explain a large proportion of variance in the Outlier dataset. The relative importance of variables associated with the Reference dataset provides the null model. Here BIO5 and BIO2 explain more variance in the Outlier SNP dataset than in the Reference dataset, which suggests that these environmental variables are important drivers of adaptive divergence.

We found that three BioClim variables explained a more rapid change in allele frequencies in the adaptive SNP datasets than in the reference datasets. The cumulative importance of allele frequency changes in the Outlier and EAA datasets differed in shape and magnitude for BIO5 (maximum temperature during the warmest month), BIO18 (precipitation during the warmest quarter), and MEM2 (Fig. 6 panels A, D, H). Most notably, the cumulative importance of the Outlier dataset showed big changes in allele frequencies at two points (19°C and 20.5°C) along BIO5 (maximum temperature during the warmest month). A similar, but slightly weaker response was seen in the EAA dataset (arrows in Fig. 6A). To examine this more closely we plotted the change in minor allele frequencies of five of the Outlier loci with the highest relative importance (R^2^) associated with BIO5 (Fig. 7A). We find that allele frequencies in R2 and R3 differ dramatically from the rest of the gradient, while frequencies in R1 are similar to R4-R6. This suggests a threshold response to the the higher temperatures experienced by R2 and R3 populations (Fig. 7B), where the adaptive genotype requires a big change in allele frequencies compared to the rest of the gradient. The rate of genomic turnover was less dramatic in response to BIO18 (precipitation in the warmest quarter, mm), but Outlier and EAA datasets both showed a higher cumulative importance than the Reference data (arrow in Fig. 6D). Finally, the Outlier dataset showed a higher cumulative importance associated with BIO2 (mean diurnal range in temperature) that was not recovered by the EAA dataset.

**Figure 6.**
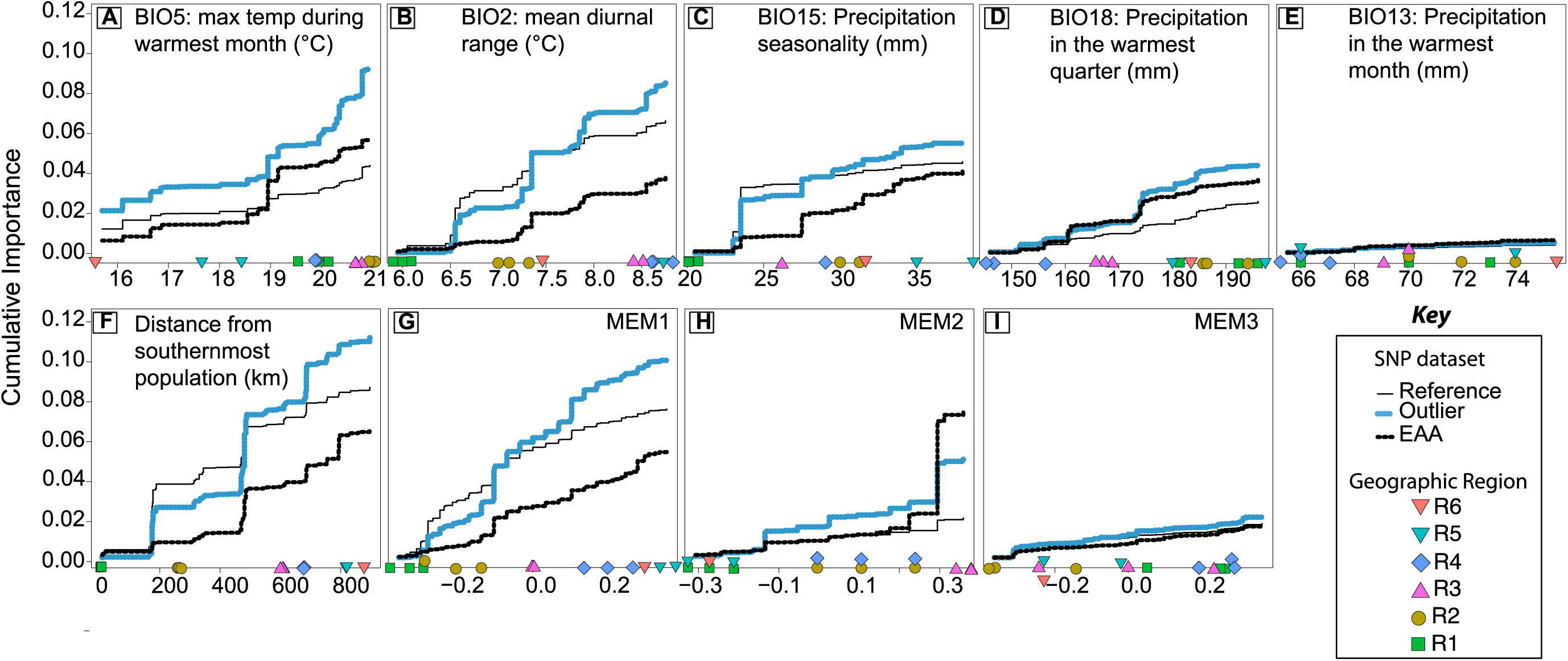
A comparison of the cumulative importance of each predictor variable for the three SNP datasets (see Key). We include the five BioClim variables (A-E) along with geographic distance (F) and three Moran Eigenvector Map variables (G-I) that explain geographic structure in the data. The maximum height of a line indicates the total allelic turnover associated with that variable. The relative importance of a particular point along the predictor variable is seen by the change in line height. The position of all populations along each variable is shown with coloured symbols along the bottom of each graph, with the populations from the same region following the same colour and shape codes as before (see Key). The Reference dataset shows the cumulative importance of each variable in explaining the change in allele frequency of neutral loci.

**Figure 7.**
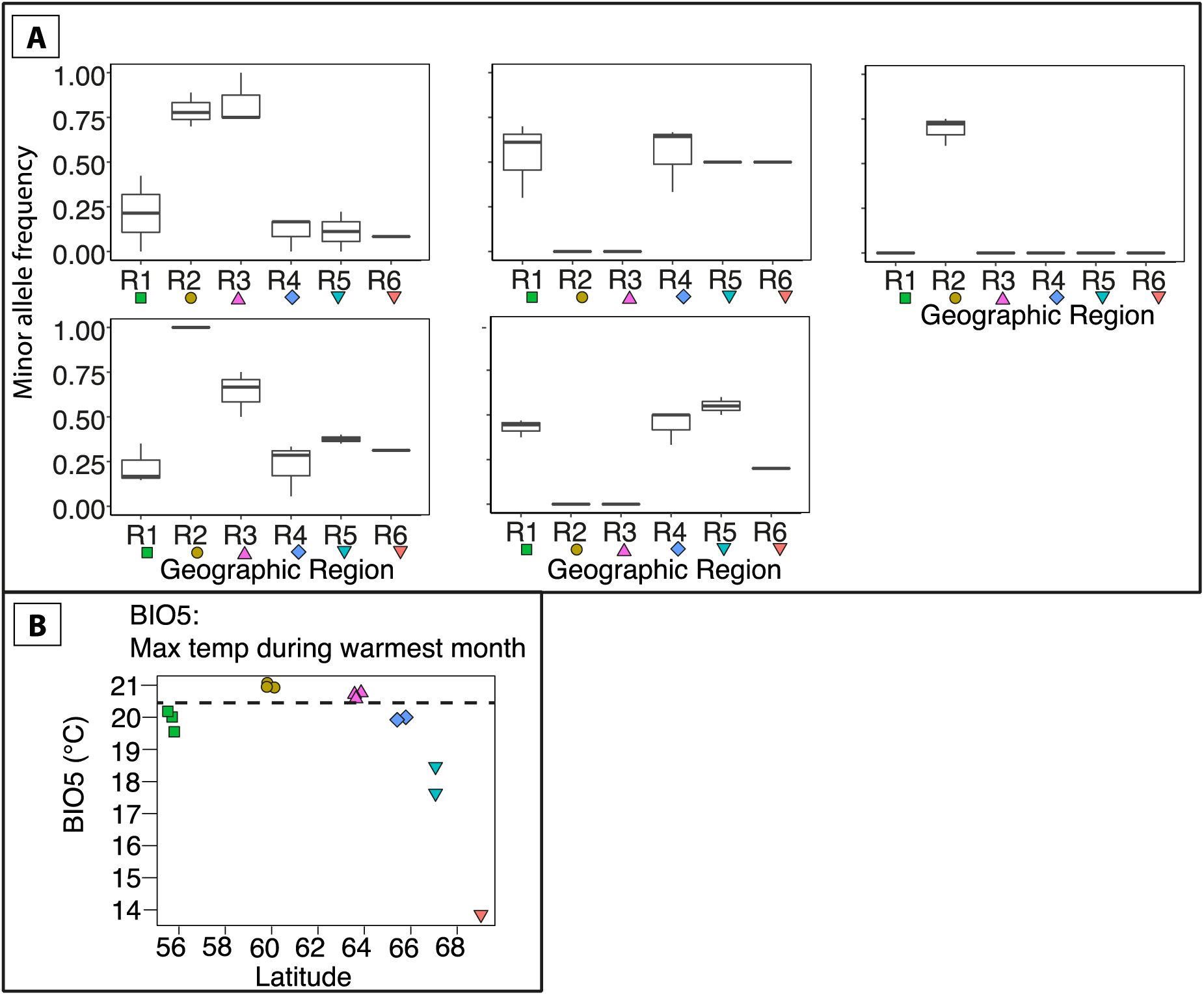
Standardised change in the minor allele frequency in five loci from the Outlier dataset that were associated with BIO5 is shown across the latitudinal gradient (A). Allele frequencies were all standardised to range between 0 and 1. Allele frequency was found to be dramatically different in R2 and R3 compared to the rest of the gradient. The change in allele frequency between R1 and R2 explains the dramatic difference between the predicted allele frequencies in the Reference and Outlier datasets (Fig. 9). The geographic distribution of BIO5 (B) is shown to illustrate the small change in temperature between R1 and R2. The dashed line indicates the temperature associated with this threshold-like response in the change in allele frequency.

Spatial mapping of the model for all three datasets showed that genomic turnover was maximal between R1 and R2, followed by the change between R2 and the rest of the populations. The three precipitation variables explained most of the variation in the Reference data allele frequencies (biplots in Fig. 8A, C, and E), while BIO5 (maximum temperature during the warmest month) was important for both the EAA and Outlier datasets. The biggest difference in genomic turnover between the adaptive loci and the Reference dataset was found in southern Sweden (Fig. 9), which suggests that the biggest change in adaptive allele frequencies is required between populations from R1 to R2.*Figure 5* The relative importance (R^2^) of the predictor variables used in the Gradient Forest analysis, calculated as the weighted mean proportion of variance in allele frequency explained by a given environmental variable. Results are shown for the five BioClim variables, geographic distance, and three Moran’s Eigenvectors (MEM1-3) explaining geographic structure in the dataset. The columns show results for the SNP datasets. EAA: Adaptive loci from environmental association analyses; Outlier: Adaptive loci identified using PCAdapt and X_T_X statistic in BayEnv2; Reference: the remaining loci. Lighter colours indicating higher importance; e.g. BIO2 and BIO5 explain a large proportion of variance in the Outlier dataset. The relative importance of variables associated with the Reference dataset provides the null model. Here BIO5 and BIO2 explain more variance in the Outlier SNP dataset than in the Reference dataset, which suggests that these environmental variables are important drivers of adaptive divergence.*Figure 6* A comparison of the cumulative importance of each predictor variable for the three SNP datasets (see Key). We include the five BioClim variables (A-E) along with geographic distance (F) and three Moran Eigenvector Map variables (G-I) that explain geographic structure in the data. The maximum height of a line indicates the total allelic turnover associated with that variable. The relative importance of a particular point along the predictor variable is seen by the change in line height. The position of all populations along each variable is shown with coloured symbols along the bottom of each graph, with the populations from the same region following the same colour and shape codes as before (see Key). The Reference dataset shows the cumulative importance of each variable in explaining the change in allele frequency of neutral loci.*Figure 7* Standardised change in the minor allele frequency in five loci from the Outlier dataset that were associated with BIO5 is shown across the latitudinal gradient (A). Allele frequencies were all standardised to range between 0 and 1. Allele frequency was found to be dramatically different in R2 and R3 compared to the rest of the gradient. The change in allele frequency between R1 and R2 explains the dramatic difference between the predicted allele frequencies in the Reference and Outlier datasets (Fig. 9). The geographic distribution of BIO5 (B) is shown to illustrate the small change in temperature between R1 and R2. The dashed line indicates the temperature associated with this threshold-like response in the change in allele frequency.

**Figure 8.**
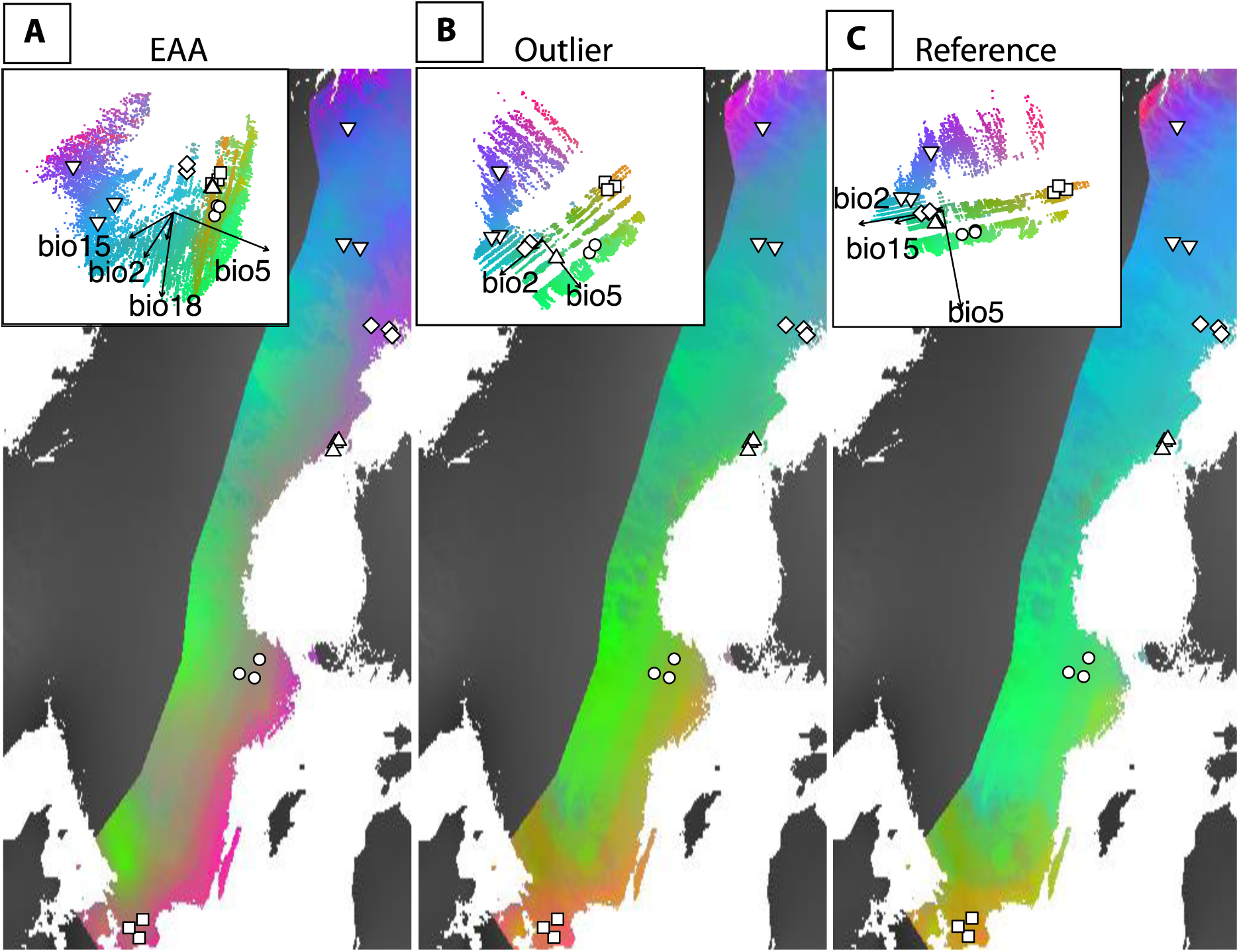
Predicted spatial distribution of genomic composition as determined for the EAA (A), Outlier (B), and Reference (C) datasets. The five BioClim variables are transformed by their relative importance in predicting genomic turnover in each dataset, and visualised as a PCA with a colour assigned to the first three principal components. Population genomic composition is expected to be similar on the same colours. The inset in each panel shows a PCA of the transformed BioClim variables, with the most important variables (see Fig. 5) shown with arrows. Sampled populations are shown on the PCA and map using the same white symbols as in Fig.1.

**Figure 9.**
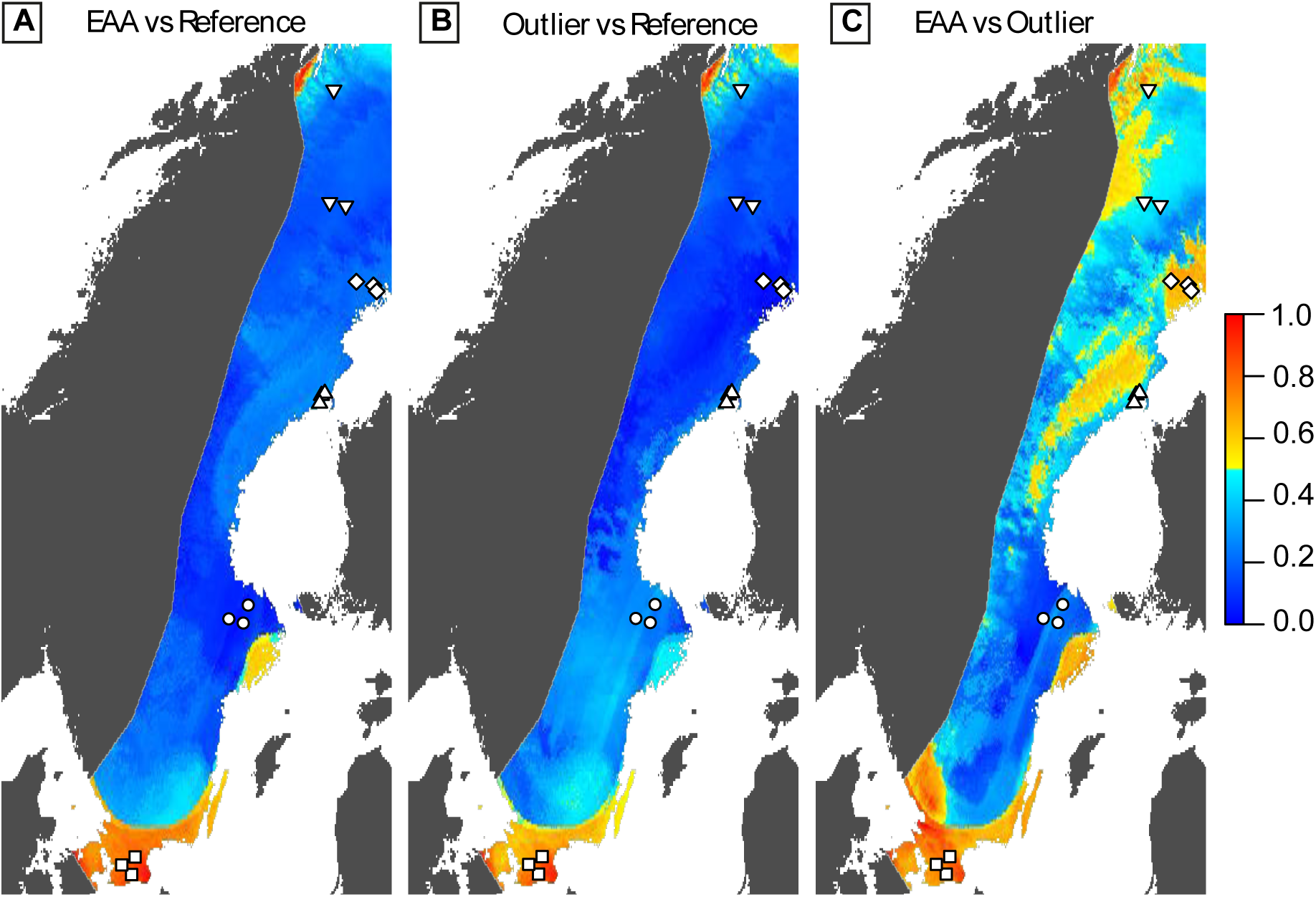
Difference in genomic turnover between the reference and adaptive datasets (a & b), and the two adaptive datasets (c). Distances were calculated as the difference between Procrustes residuals in the matrix comparisons and scaled by the maximum distance found for each comparison. Large differences between datasets are indicated with warmer colours. Sampled populations are shown using white symbols (see Fig. 1).

## Discussion

By combining landscape genetics and genomic turnover analyses we describe how climate and geography structure *R. temporaria* genomic variation across the Scandinavian latitudinal gradient. Genomic variation in this system is strongly related to geography, but also shows evidence of adaptation to climate. A surprisingly large portion of the total genomic variation is attributable to climate variables (38%) or geographically structured climate (30%). We also find a threshold response in to BIO5 (maximum temperature in the warmest month), implying that a thermal threshold occurs in southern Sweden. Finally, our results show that the biggest mismatch between neutral and adaptive allele frequencies occurs in southern Sweden, largely driven by the threhold response to BIO5. Our results show that an analysis of the geographic distribution of genomic variation in *R. temporaria* provide important insights into the climatic drivers and potential adaptive thresholds across a well-studies system.

### Population Structure

We found strong population structure across the latitudinal gradient, with the biggest divergence in allele frequencies between R1 and the rest of Sweden. Sequential pairing off of the rest of the populations and strong signals of IBD are indicative of an expansion from a southern colonisation. These results support previous work based on mtDNA that has suggested a single colonisation route and northward expansion in Scandinavia by *R. temporaria* (Palo *et al*. 2004). Previous phylogenetic work from the same study indicated that all populations in Scandinavia and northwards assign to the eastern mitochondrial haplogroup (Palo *et al*. 2004), and the contact zone between the eastern and western lineages has been described in Northern Germany (Schmeller *et al*. 2008). However, population assignment analysis based on microsatellite data found that some southern Scandinavian populations (including from R1) assigned up to 100% to the Western lineage based on multilocus genotypes (Palo *et al*. 2004). This explains the divergence of the southern populations, perhaps even up to R2, from the rest of the gradient. Overall our results support previous work that asserted that two mitochondrial lineages colonised Scandinavia from the south; via Denmark and Sweden. A contact zone between these lineages in southern Sweden resulted in the divergent allele frequencies in this region compared to the rest of the gradient. The eastern mitochondrial haplotype occur throughout extant Scandinavian populations, which suggests that gene flow from the populations with the eastern haplogroup have since swamped and replaced the western haplogroup (Palo *et al*. 2004).

### Geography and Climate determine genotype

We found that climate and geographically structured climate explained a large proportion (76.6%) of the total genomic variation. Climate independent of geography and geographically structured climate explained similar amounts of variation (38% and 30%, respectively). Strong clinal genetic structure across the Scandinavian latitudinal gradient which has been attributed to consistent selection gradients co-varying with geography (Palo *et al*. 2003b, 2004; Cano *et al*. 2004). However, our results suggest that a significant proportion of the adaptive divergence across the gradient could be associated with climate variables that are not latitudinally ordered. Season length (number of days above 6 °C, the developmental threshold for *R. temporaria* tadpoles; Laurila *et al*. 2001; Laugen *et al*. 2003b; Muir *et al*. 2014a) and temperature during larval development (30 days after spawning) are two environmental variables that are commonly attributed to the latitudinal adaptive divergence (e.g. Laugen *et al*. 2003b, 2005a). While season length is latitudinally ordered, water temperatures during the larval phase peak at mid-latitudes (Laugen *et al*. 2003b). Common garden experiments have found that several larval traits - egg development time, size at hatching, larval growth rate, size at metamorphosis, and resting metabolic rate - follow this curvilinear distribution across latitude (Pahkala *et al*. 2002; Laugen *et al*. 2003b, 2005b; Palo *et al*. 2003b; Lindgren & Laurila 2005, 2009). Adult body size, skeletal growth, and lifetime activity follow the same curvilinear distribution, and are maximised at mid-latitudes (Laugen *et al*. 2005b; Hjernquist *et al*. 2012).

Together with our results show support for latitudinally ordered adaptive divergence, but also present evidence that climatic drivers that are not latitudinally ordered are important for adaptation.

### Non-linear changes in allele frequencies and threshold effects

We find a strong threshold effect in a subset of Outlier loci associated with BIO5 (maximum temperature during the warmest month). This suggests that there is a physiological threshold in response to BIO5 (or something related to BIO5). Thesholds in polygenic traits are likely to be common in heterogenous environments (Roff 1996). Indeed, we find that four of the five BioClim variables showed an elevated cumulative importance for the adaptive loci compared with the Reference datasets. Of these, BIO2 (mean diurnal range in temperature) and BIO18 (precipitation during the warmest quarter) show evidence of thresholds at which allele frequencies change more rapidly than in the Reference dataset. Geographically the BIO5 threshold separates R2 and R3 from the rest of the gradient. We found dramatically different allele frequencies in a set of Outlier loci associated with BIO5 in this region. Comparison in genomic turnover between the datasets identified the transition between R1 and R2 to diverge the most between the Reference and adaptive datasets. This is indicative of strong adaptive divergence in this region.

Adaptive divergence across the latitudinal gradient in Europe has been extensively documented in plants and animals (e.g. Laugen *et al*. 2003b; Debieu *et al*. 2013; Vergeer & Kunin 2013). Much of this work is based on common garden and reciprocal transplant experiments have established divergence in various phenotypes across the environmental gradient. Landscape and population genomic methods provide a powerful approach to complement and extend these results in several ways. These include determining the proportion of genomic variation associated with adaptation, identifying the genomic underpinnings and architecture of adaptation, identifying important climatic drivers of adaptive divergence, and identifying adaptive thresholds in response to a specific variable. More generally, this approach can have valuable conservation implications, particularly for mitigating the loss of biodiversity due to climate change. One of the most valuable outcomes lies in identifying populations where a small change in environment will result in a large mismatch between genotype and environment. These populations are particularly vulnerable to extinction, and conservation management action would have to be carefully considered.

